# PharmacoDB 2.0 : Improving scalability and transparency of *in vitro* pharmacogenomics analysis

**DOI:** 10.1101/2021.09.21.461211

**Authors:** Nikta Feizi, Sisira Kadambat Nair, Petr Smirnov, Gangesh Beri, Christopher Eeles, Parinaz Nasr Esfahani, Minoru Nakano, Denis Tkachuk, Anthony Mammoliti, Evgeniya Gorobets, Arvind Singh Mer, Eva Lin, Yihong Yu, Scott Martin, Marc Hafner, Benjamin Haibe-Kains

## Abstract

Cancer pharmacogenomics studies provide valuable insights into disease progression and associations between genomic features and drug response. PharmacoDB integrates multiple cancer pharmacogenomics datasets profiling approved and investigational drugs across cell lines from diverse tissue types. The web-application enables users to efficiently navigate across datasets, view and compare drug dose-response data for a specific drug-cell line pair. In the new version of PharmacoDB (version 2.0, https://pharmacodb.ca/), we present: (*i*) new datasets such as NCI-60, the Profiling Relative Inhibition Simultaneously in Mixtures (PRISM) dataset, as well as updated data from the Genomics of Drug Sensitivity in Cancer (GDSC) and the Genentech Cell Line Screening Initiative (gCSI); (*ii*) implementation of FAIR data pipelines using ORCESTRA and PharmacoDI; (*iii*) enhancements to drug response analysis such as tissue distribution of dose-response metrics and biomarker analysis; (*iv*) improved connectivity to drug and cell line databases in the community. The web interface has been rewritten using a modern technology stack to ensure scalability and standardization to accommodate growing pharmacogenomics datasets. PharmacoDB 2.0 is a valuable tool for mining pharmacogenomics datasets, comparing and assessing drug response phenotypes of cancer models.

**HIGHLIGHTS:**

- PharmacoDB 2.0 includes new and updated large pharmacogenomic datasets. The data processing for PharmacoDB is made fully reproducible through the use of the ORCESTRA platform and automated data ingestion pipelines
- The new release contains enriched annotations for drugs and cell lines via connectivity to external databases, as well as new analytical methods for tissue-specific and pan-cancer biomarker discovery
- The new version of PharmacoDB incorporates a scalable and reproducible framework that can accelerate the implementation of analytical pipelines including machine learning/AI for biomarker discovery in the future

## INTRODUCTION

With the advent of high-throughput technologies, a vast amount of genomic data is being generated across various disease domains. In oncology, genomic and pharmacological profiling of cancer cell line models has resulted in a better understanding of the relationship between the molecular features of cancers and treatment outcomes. Starting with a disease-oriented screening model in the late 1980s, the US National Cancer Institute’s NCI-60 anticancer drug screen has aided in major discoveries across many fields including anticancer therapy (1). Subsequently, several studies, including the Genomics of Drug Sensitivity in Cancer (GDSC) (2, 3), Cancer Therapeutics Response Portal (CTRP) (4), and Cancer Cell Line Encyclopedia (CCLE) (5), have generated pharmacogenomic profiles of much larger panels of cancer cell lines. Collectively, such data can be used for hypothesis testing and in the discovery or repurposing of new anti-cancer therapeutics.

PharmacoDB (version 1.0) was released in 2018 (6) as the largest database integrating cancer cell line pharmacogenomic datasets, enabling users to efficiently explore data across the largest published studies. PharmacoDB provided a unified analysis platform by standardizing statistical models of dose response data, and harmonizing annotations of experiments. The next generation of PharmacoDB (version 2.0) includes new datasets such as the US National Cancer Institute’s NCI-60 (7–12), the Broad’s Profiling Relative Inhibition Simultaneously in Mixtures (PRISM) dataset (13), GDSC version 2 (GDSC2), and updates to existing datasets such as the Wellcome Trust Sanger Institute’s GDSC version 1 (GDSC1) (14) and the Genentech Cell line Screening Initiative (gCSI) (15) (Table 1). The cell lines investigated in these studies were screened for drug response and where available, profiled at the molecular level with multiple technologies, including RNA sequencing, microarray single-nucleotide and gene expression profiling, and whole-exome or whole-genome sequencing. The processing of the pharmacogenomic data is fully automated and documented to generate FAIR (Findability, Accessibility, Interoperability, and Reusability) data through the use of ORCESTRA (https://orcestra.ca/) and PharmacoDI ingestion pipelines. PharmacoDB 2.0 also provides new visualization of differential drug dose-response across tissues, as well as summaries of gene-drug associations showcasing their strength and reproducibility across studies. New links to drugs and cell lines have been added from Reactome (16), Drug Target Commons (DTC) (17) and Cellosaurus (18) to increase the connection to PharmacoDB from other databases in the community. The chemical identifiers are extended to include ChEMBL (19) IDs (Figure 1A).

**Table 1.**
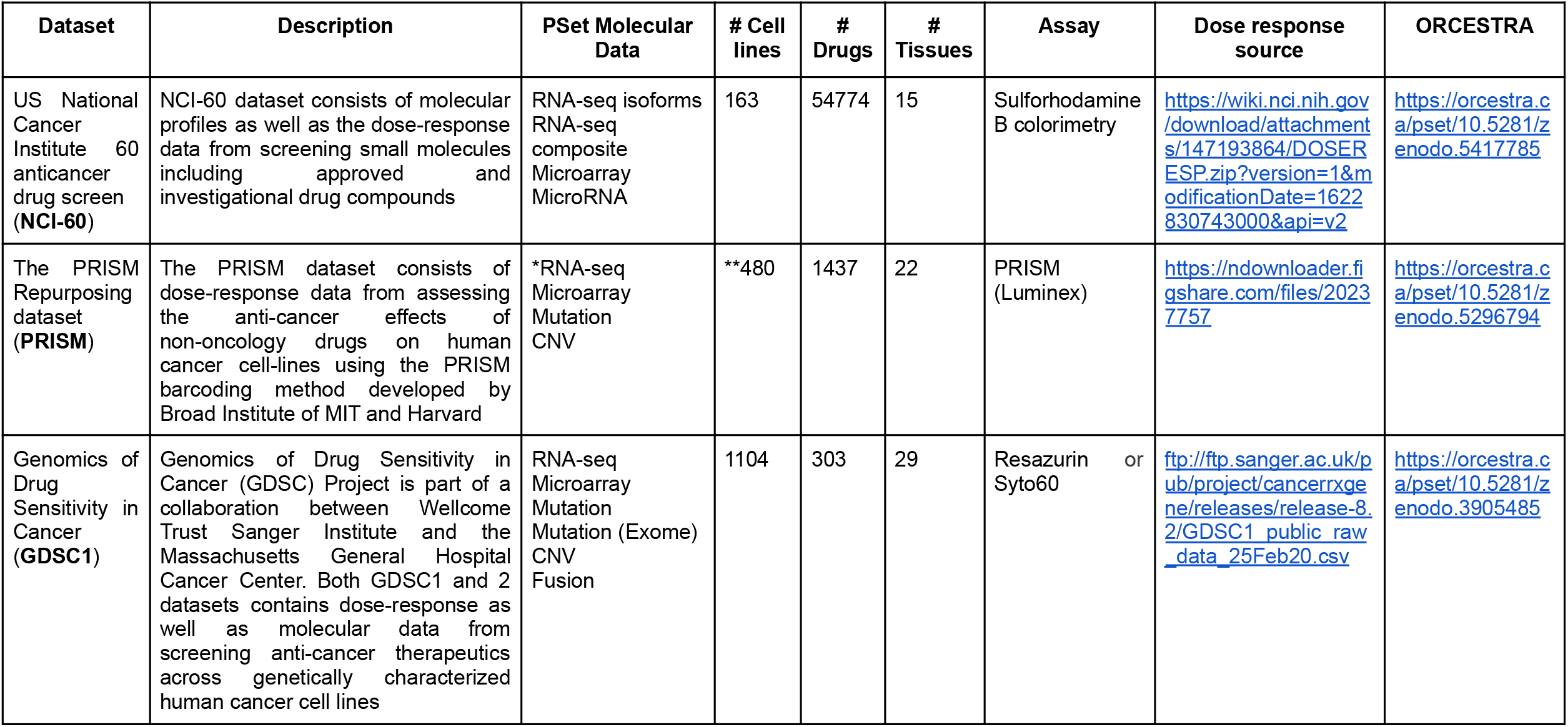
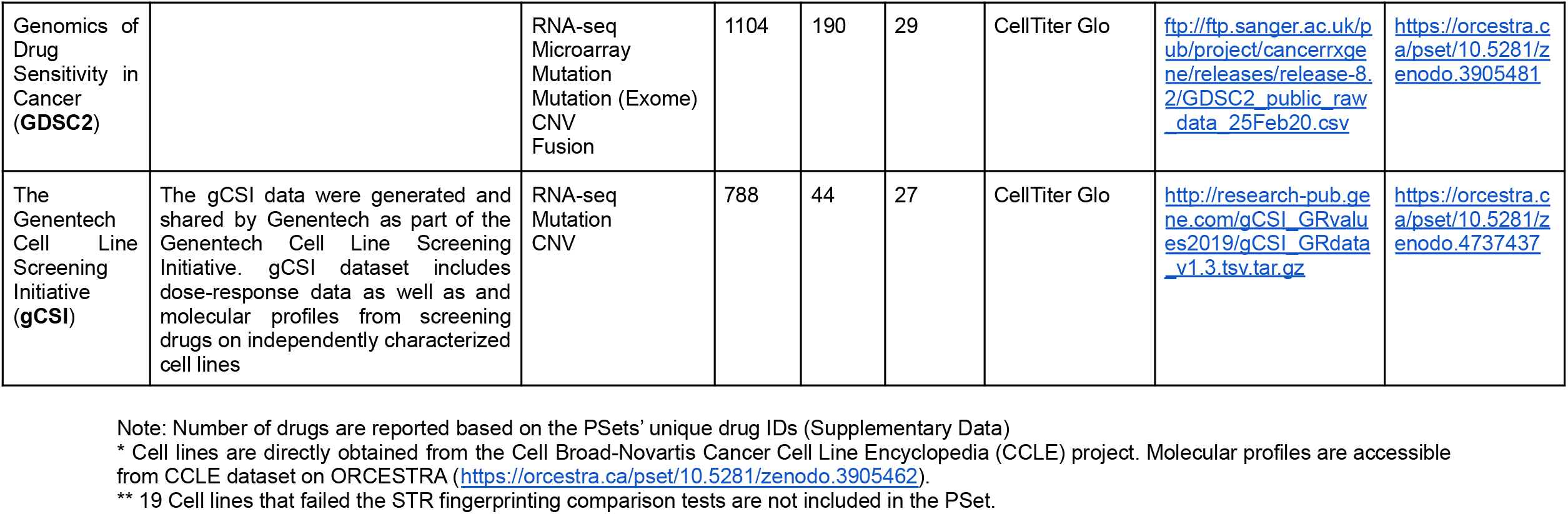
Details of new and updated datasets in PharmacoDB 2.0. An overview of the new and updated datasets, types of molecular profiles included in each dataset, number of cell lines, drugs, and tissue types in each dataset, and the assay type used for measuring the dose response values. The link to both the source of raw dose response data and the corresponding PSets on ORCESTRA is provided in the table.

**Figure 1 -.**
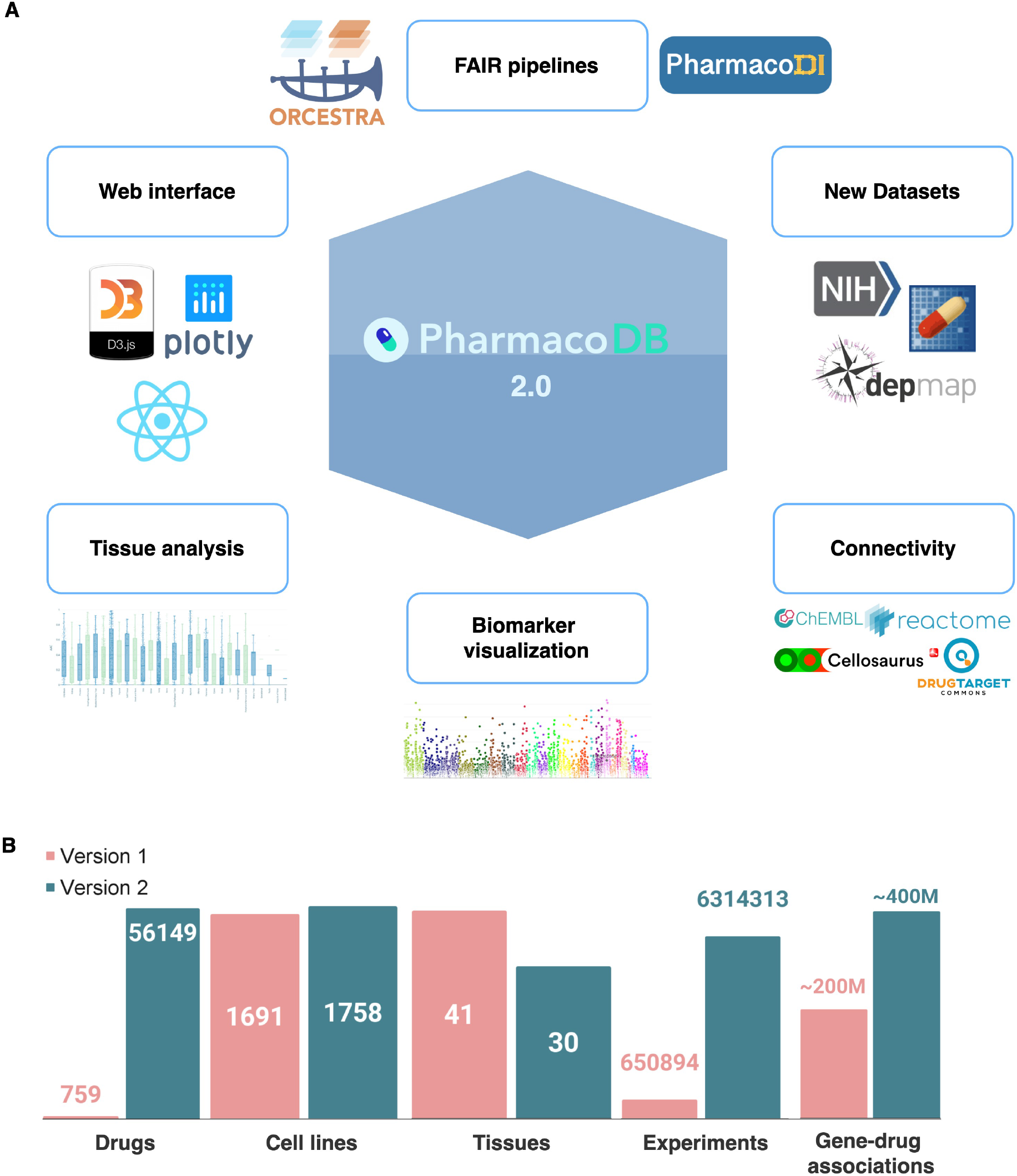
PharmacoDB 2.0 overview. **A)** The new version of PharmacoDB includes updated and new large-scale pharmacogenomic datasets. The web-application contains enriched annotations for drugs and cell lines via connectivity to external databases. PharmacoDB 2.0 includes new analytical methods for tissue-specific and pan-cancer biomarker discovery. The new web-interface ensures scalability and simplifies maintenance. PharmacoDB 2.0 is made fully reproducible through the use of the ORCESTRA platform and automated data ingestion pipelines. **B)** Bar plots showing previous (Version 1) and current (Version 2) database statistics.

## ADDITION OF NEW DATASETS

The current release of PharmacoDB includes significant updates to the GDSC1000 (renamed GDSC1 following the Wellcome Trust Sanger Institute’s nomenclature) and gCSI datasets (Table 1). In addition, the NCI-60, PRISM, and GDSC2 datasets have been included in this new release of PharmacoDB. The numbers of entities such as drugs, cell lines, tissues, gene-drug associations and experiments in PharmacoDB 2.0 and the previous version is shown in (Figure 1B). The newly standardized tissue curation is explained in Supplementary Data.

The NCI-60 screen includes over 55,000 small molecules assayed on 60 core (and 103 additional) cell lines representing 15 tumor types. PharmacoDB 2.0 includes percentage of treated cell growth (PTC) values, downloaded from the NCI Developmental Therapeutics Program (https://wiki.nci.nih.gov/display/NCIDTPdata/NCI-60+Growth+Inhibition+Data), for 4,557,787 experiments that included at least 4 measured doses required for curve fitting (Supplementary Data). The PRISM Repurposing dataset employs a molecular barcoding method to screen drugs against cell lines in pools (13, 20). The barcoded cell lines from different lineages are assayed for relative mRNA abundance after treatment with a drug or chemical perturbagen to estimate cell viability. PRISM drug screening involves a 2-stage screening strategy: a) Screen-1 includes 4518 compounds and 578 cell lines that were assayed in triplicate at a single dose and b) Screen-2 includes 1,448 drugs that were re-screened against 499 cell lines in triplicate in an 8-point dose-response (13). PharmacoDB 2.0 includes the Screen-2 dose-response data with biological replicate-collapsed log-fold change values, downloaded from the Dependency Map Data Portal (https://depmap.org/repurposing/). After processing, these data included 726,814 experiments with at least 4 dose measurements (Supplementary Data).

The GDSC2 dataset was generated using a new screening platform from the Wellcome Trust Sanger Institute’s Genomics of Drug Sensitivity in Cancer project (21, 22). The dataset, previously included as GDSC1000 in PharmacoDB, has been renamed to GDSC1 and updated to include more experiments (323,032 GDSC1 *vs*. 225,480 GDSC1000), reflecting the updated data available from the GDSC project. The GDSC2 dataset represents a different screening approach (using the same cellular viability assay as the Broad and Genentech studies, and an increase in biological replicates) on a similar set of cell lines as GDSC1, with both overlapping and new compounds, and a total of 215,780 experiments which were included in PharmacoDB 2.0. More information on the protocol differences between the GDSC1 and GDSC2 data can be found on the GDSC website at https://www.cancerrxgene.org/. Finally, PharmacoDB 2.0 includes significant updates to Genentech’s gCSI dataset. The number of experiments available has increased from 6,455 to 16,688, primarily through the inclusion of additional 35 compounds and extensive biological replicate information.

For all new additions and updates to the database, drug annotations were curated by mapping drugs to PubChem (23) identifiers such as compound identifiers (CID), SMILES, and InChIKeys using the PUG-REST API (24). Whenever possible, PubChem index names were used as the standard name in PharmacoDB. Most exceptions occurred in the NCI-60 dataset, where a large proportion of PubChem index names followed IUPAC nomenclature and were difficult for humans to read. For these, we followed the process described in Supplementary Figure S1 to decide when to prefer the NCI-60 dataset ID as the standard name for PharmacoDB. Cell lines were annotated using Cellosaurus (18), an online cell line knowledge resource which documents cell lines used in biomedical research. OncoTree (25) was used to re-label the tissues of cancer cell lines profiled into an externally defined ontology.

## IMPLEMENTATION OF REPRODUCIBLE PIPELINES

With large amounts of pharmacogenomic data released from multiple studies, the reproducibility of computational pipelines used to process these multimodal data is essential. This includes adhering to the FAIR data principles, along with ensuring that there is a standardized manner in which the pipelines are executed and the data is hosted. To address this important issue, we used ORCESTRA (https://orcestra.ca/), a platform that allows researchers to process biomedical data into unified data objects in a reproducible and transparent manner, where data provenance is tracked (26). At the heart of ORCESTRA is Pachyderm (https://www.pachyderm.com/), an open-source data versioning tool used to execute pipelines processing the molecular and compound screening data, and packaging the datasets into R objects called PharmacoSets (PSets), which are implemented by the PharmacoGx package (27). At the end of each PSet processing pipeline, the created object is automatically deposited on Zenodo and assigned a digital object identifier (DOI). Once the highly curated and standardized PSets are released via ORCESTRA, they need to be preprocessed into tables which match the PharmacoDB Entity Relationship Diagram (ERD) before being loaded into the database. To ensure that the data ingestion standards in PharmacoDB adhere to FAIR data principles, the Pharmaco-Data Ingestion (PharmacoDI) project was initiated to create an Extract Transform Load (ETL) pipeline which adheres to modern data engineering best practices (28, 29).

The PharmacoDI project consists of three major components. Firstly, the rPharmacoDI (https://github.com/bhklab/rPharmacoDI) R package provides an interface to download and export PSets in a Python compatible format. Secondly, the PharmacoDI (https://pypi.org/project/PharmacoDI) Python package contains a set of functions for transforming the exported raw PSet data into tables which match the PharmacoDB schema, leveraging the Python Datatable package to allow larger than memory data processing. Finally, the Snakemake workflow management tool is used to integrate ORCESTRA, rPharmacoDI, PharmacoDI and PharmacoDB into a fully modular and scalable ETL pipeline (https://github.com/bhklab/PharmacoDI_snakemake_pipeline) which ensures that all software dependencies, metadata and code are version controlled, transparent and fully reproducible. This pipeline includes a number of additional features which keeps PharmacoDB annotations up to date, such as dynamic queries to ChEMBL (19) and Cellosaurus (18) to fetch respective drug and cell line metadata. Automated quality control checks are implemented throughout the pipeline to ensure data integrity and flag data for manual review if problems are detected. Once quality control passes, the Python SQLAlchemy package is used to connect with our Azure MySQL database where database tables are automatically created and loaded before being deployed for use in the PharmacoDB web-application (Figure 2).

**Figure 2 -.**
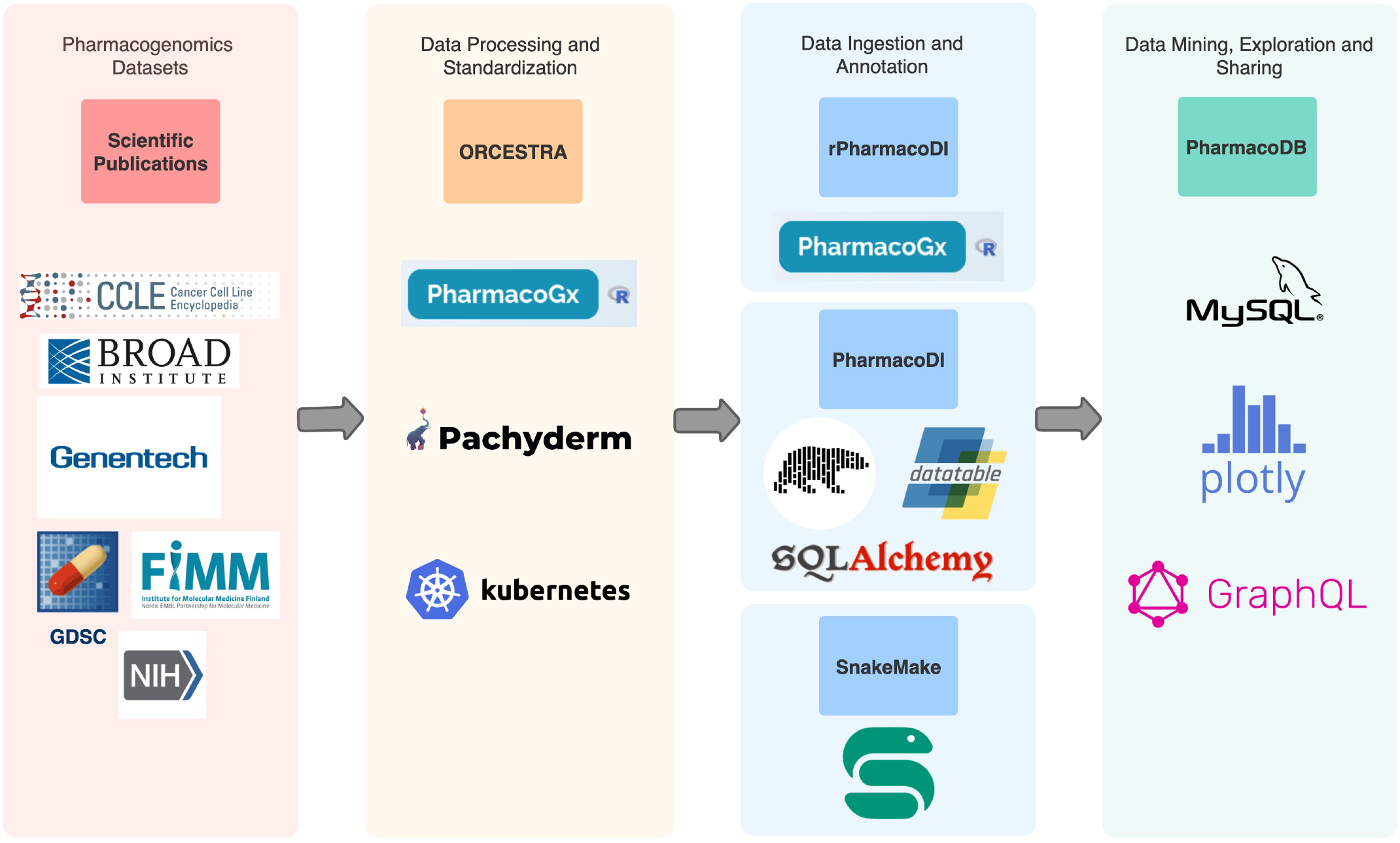
Computational processing pipeline of raw pharmacogenomic data for ingestion into PharmacoDB. Different panels show the process of ingesting public datasets into PharmacoDB 2.0. The first panel highlights the sources of the newly added datasets, while the subsequent panels highlight the tools and technologies used for Data Processing and Standardization, Data Ingestion and Annotation, and for building the PharmacoDB 2.0 web app itself.

## CONNECTIVITY TO EXTERNAL DATABASES

While the original PharmacoDB database included links to external databases for drugs, genes, and cell lines, PharmacoDB 2.0 focuses on creating bi-directional links to improve discoverability of the available pharmacogenomic data. Unique stable identifiers were created for both drugs and cell lines in the PharmacoDB database, allowing external databases to link directly to entities in PharmacoDB. A collection of drugs from PharmacoDB 2.0 are bi-directionally linked to Reactome (16), an online database of biological pathways including drug mechanisms of action. Reactome provides detailed insights into drug targets, binding partners, and subsequent biological pathways associated with target action. PharmacoDB drugs are also linked to Drug Target Commons (DTC) (17), a community-driven web platform for compound-target bioactivity assay annotation profiles relevant for drug discovery and repurposing. ChEMBL (19) IDs matching our compound identifiers are added in addition to PubChem identifiers. The cell lines from the datasets are bi-directionally linked and annotated to Cellosaurus (18), which is a cell line information resource. Additional cell line metadata such as disease, metastasis site, species, and links to external databases such as DepMap are available from Cellosaurus for the linked cell lines. The external links can be found in the individual drug and cell line pages of PharmacoDB as well as in external web-applications. In addition, PharmacoDB APIs are implemented using JavaScript programming language in Express which is a back end web application framework for NodeJS. GraphQL (https://graphql.org/), which is an open-source data query and manipulation language for the APIs is used to structure the APIs, and which also provides a runtime for fulfilling queries with existing data. The newly created APIs provide better performance and are open source to facilitate user operability. The APIs can be accessed without any authentication process or tokens.

## ENHANCED DATA ANALYSIS

### Tissue-specific analysis

Building on top of the newly standardized tissue ontology, PharmacoDB 2.0 has been updated with a focus on tissue-specific analysis throughout the web application. PharmacoDB 2.0 contains cell response data for a large panel of 1758 cell lines spanning across 30 tissues. Moreover, PharmacoDB includes data for a large portfolio of 589 FDA approved drugs, several investigational drugs, tool or lead compounds, and natural substances, all with varying levels of activity across and within tissue types. Therefore, the visualizations within the web-application have been extended to help identify patterns of sensitivity across and within tissue(s). On each individual drug page, the differential sensitivity to a compound across tissues is displayed as a boxplot, giving a quick overview of the tissues that are sensitive or resistant to a compound. As with all the plots in the web-application, this can be displayed by integrating data across all datasets, or filtered by dataset(s). For example, viewing this plot across all datasets for Dabrafenib clearly indicates the well known preferential sensitivity of BRAF mutated skin cancers to BRAF inhibition (30), and its absence in other tissues (even tissues known to harbour mutations in the BRAF pathway, such as bowel) (Figure 3A). Alternatively, within PharmacoDB 2.0 it is possible to investigate the differential sensitivity to a compound within a tissue type by comparing the drug dose response curves across all cell lines tested in all datasets for a particular tissue. Continuing with the example of Dabrafenib, it is possible to identify the most sensitive and resistant skin cell lines, using either the sensitivity summary metrics provided in PharmacoDB (such as the Area Above the Curve - AAC or the drug concentration necessary to inhibit 50% of the maximal cell viability - IC50), or visually (Figure 3B). This will be a useful addition for experimentalists interested in identifying the most sensitive and/or resistant models to a particular compound for investigating mechanism, synthetic lethality, or possible drug combinations.

**Figure 3 -.**
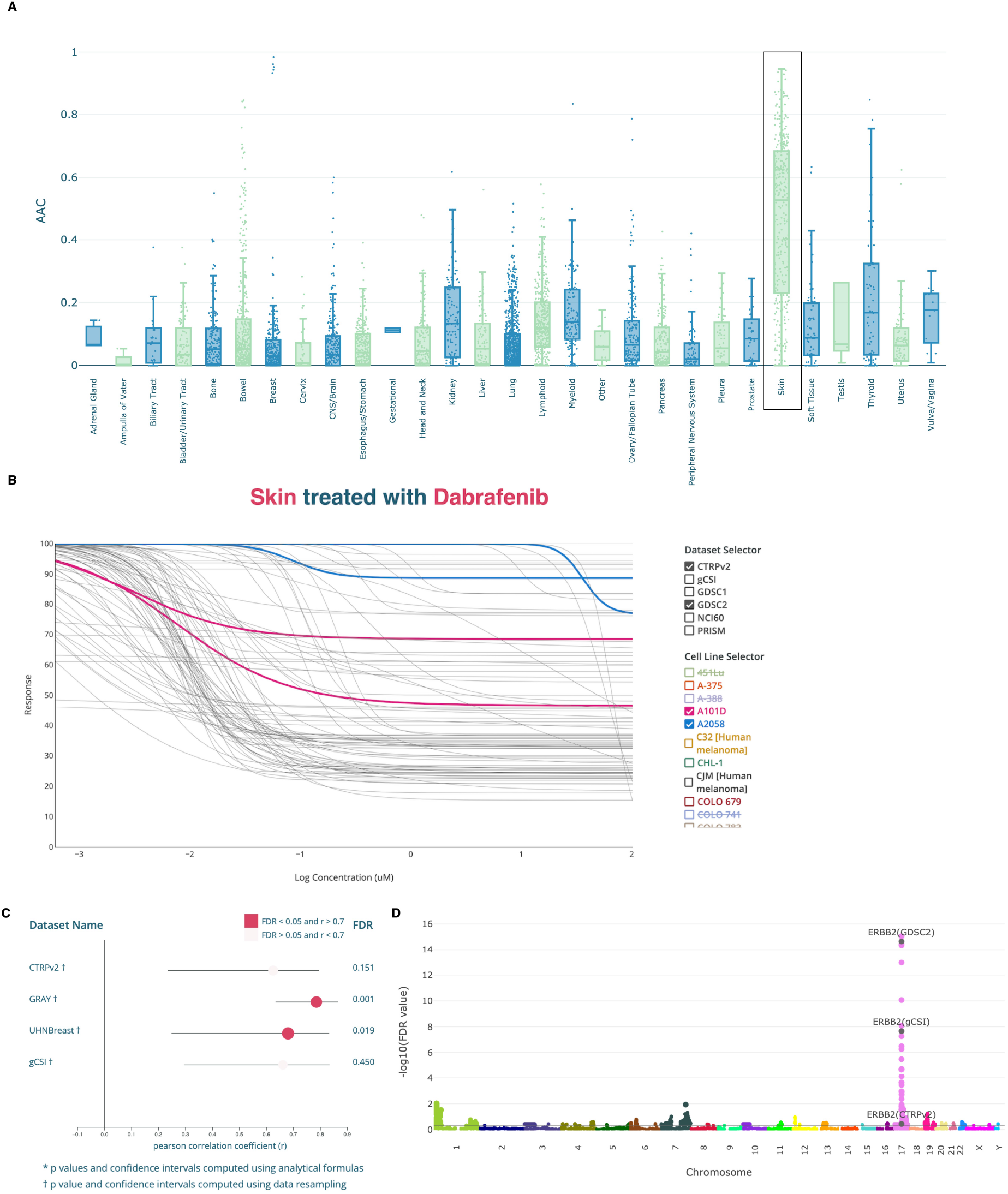
Visualization of tissue-specific drug response and gene-drug associations. **A)** Drug response (AAC) of Dabrafenib across various tissues from all datasets. **B)** Differential sensitivity of skin cell lines to Dabrafenib; cell lines and datasets of interest can be highlighted in the plot by checking the boxes. **C)** Forest plot of Pearson correlations between Lapatinib response and ERBB2 expression in breast tissue. Data from rna sequencing is shown here. The significant associations (FDR < 0.05 and pearson correlation coefficient, r > 0.7) is highlighted in bright pink. **D)** Manhattan plot showing the association of copy number alterations with Lapatinib response in all datasets and across all tissue types, with ERBB2 highlighted. The genomic coordinates are displayed on the x-axis, and negative logarithm of the association p-value is displayed on the y-axis. The different colors of each block show the extent of each chromosome.

### Query biomarkers

PharmacoDB 2.0 improves the users’ ability to explore, evaluate, and compare the molecular features most strongly correlating with response to particular compounds across pharmacogenomic studies. The original version of PharmacoDB contained precomputed tables for pan-cancer associations of drug response with gene expression, mutation, and copy number variation computed using the *PharmacoGx* package (27). The data were presented in a plain table format, useful only for ranking associations by significance or effect size. PharmacoDB 2.0 extends this by providing interpretable visualizations to compare and contextualize these associations, extends the analysis to tissue-specific associations, and incorporates updates to the statistical methodology to evaluate the strength and significance of these markers.

The previous pipeline used to evaluate the associations between all feature types, and drug response was based around a linear modelling approach, and relied on analytical p-values derived from assumptions of normality. However, both the distributions of molecular features and drug response metrics can deviate significantly from normal, potentially inducing biases in the estimation of these p-values (31). This is exacerbated by our addition of tissue-specific associations to the database, for which sample sizes are much smaller (dozens vs. hundreds of cell lines). Therefore PharmacoDB 2.0 includes both analytic and permutation-estimated p-values for associations, increasing the reliability of statistical significance estimates in the database (details on statistical methodology are available in the Supplementary Data).

Tissue-specific gene-compound associations require evaluating the correlation between a feature and drug response within cell lines limited to a single tissue type. Repeating this for each of the 30 tissues in PharmacoDB becomes a significant undertaking, increasing the table associations from approximately 33.5 million pan-cancer associations to over 400 million tissue specific associations (after filtering out associations based on less than 20 cell lines). While analytical test results are available for all associations, permutation testing of all 400 million associations is a lengthy and ongoing process. Currently, PharmacoDB contains the results of permutation testing associations appearing in at least 3 datasets. As we compute the results of permutation tests for more compounds and gene features, we will be continuously updating the database.

Finally, each gene-compound association (tissue-specific or pan-cancer) is now associated with a biomarker page within the web application, which contains summary information about the gene, the compound, and two types of plots contextualizing the associations available for this gene and compound pair in the database. A forest plot allows users to compare the strength of associations between the compound and gene across datasets. In Figure 3C, we show the association between the validated biomarker of ERBB2 expression (32, 33) and Lapatinib response in breast tissue. The forest plot gives a visual summary of the reproducibility of this marker across studies. As a forest plot is displayed for each available molecular feature type, it also facilitates comparison of the drug response effect between expression, copy number, and mutation alterations of the same gene. The second plot, specifically designed for copy number alteration (CNA) associations, is a Manhattan plot, displaying the strength and significance of each CNA association with this particular compound across the dataset (Figure 3D). This provides important context to the association, allowing a visual identification of associations that are driven by focal copy number changes, versus those that are likely passengers to larger genomic events.

## CONCLUSION AND FUTURE DIRECTIONS

Large-scale compound screens across various biological model systems are being carried out at a fast pace, generating valuable pharmacogenomic data for biomarker discovery, a key challenge in precision medicine. PharmacoDB 2.0 is a major update bringing enhancements to the User Interface of the web application, greatly expands the pharmacogenomic data available within the database, and implements pipelines following FAIR principles which allow researchers to track the provenance of data included in each release of PharmacoDB going forward. PharmacoDB provides a platform of reference to cancer researchers while designing their experiments. It can be used either for checking if an experiment has been carried out by other research groups or to compare experiment outcomes with public data. Information on the cell line and tissue breakdown of drug studies in datasets, sensitivity across tissues, and top gene-drug associations are particularly helpful in this regard. PharmacoDB also helps users check the association of their gene of interest with drug response and further analyze the reliability of a potential biomarker across various studies. Finally, PharmacoDB serves as a tool for drug repurposing by providing drug response on tissue types other than the drug’s approved indication.

Since the first publication of PharmacoDB in 2018, several other web applications have arisen which integrate pharmacogenomics studies, including CellMinerCDB (34), and the Dependency Map Portal integrating GDSC, CTRP, and PRISM drug response data with RNAi and CRISPR essentiality screening data (3, 4, 13, 20). While these tools have overlapping uses, PharmacoDB has and continues to focus on integrative and comparative analysis across a wide range of published pharmacogenomics studies, and as such includes major pan-cancer screening initiatives as well as smaller, tissue-specific studies. Neither CellMinerCDB nor the Dependency Map portal integrate the smaller pan-cancer gCSI dataset or the breast cancer specific GRAY and UHNBreast datasets.

Future planned work includes a focus on including an increasing number of smaller published screens as well as the integration of formal statistical meta-analysis within the web application to leverage the growing number of studies for biomarker discovery. PharmacoDB has already proven to be a valuable resource for computational and wet-lab researchers and with continued improvements, we aim to make pharmacogenomics data more discoverable and accessible for the cancer research community.

## DATA AVAILABILITY

The Pharmacosets can be downloaded from https://orcestra.ca/, with the link to each dataset being provided in Table 1.

## CODE AVAILABILITY

The code for PharmacoDB development is available on https://github.com/bhklab/PharmacoDB-JS. Code for rPharmacoDI and PharmacoDI python packages are https://github.com/bhklab/rPharmacoDI and https://github.com/bhklab/PharmacoDI respectively. SnakeMake database creation pipelines can be found here - https://github.com/bhklab/PharmacoDI_snakemake_pipeline

## SUPPLEMENTARY DATA

Supplementary Data are available at NAR Online.

## ACKNOWLEDGMENT

We thank Richard Bourgon for his thoughtful suggestions.

## FUNDING

This work was supported by Genome Canada Bioinformatics and Computational Biology [15414]; Princess Margaret Data Science Program; Ontario Institute for Cancer Research (OICR; PanCuRx Translational Research Initiative) through funding provided by the Government of Ontario. Funding for open access charge: Genome Canada.

## CONFLICT OF INTEREST

No conflict of interest to declare.

## Notes

### Competing Interest Statement

The authors have declared no competing interest.

## REFERENCES

1. Shoemaker, R.H. (2006) The NCI60 human tumour cell line anticancer drug screen. Nat. Rev. Cancer, 6, 813–823.

2. Garnett, M.J., Edelman, E.J., Heidorn, S.J., Greenman, C.D., Dastur, A., Lau, K.W., Greninger, P., Thompson, I.R., Luo, X., Soares, J., et al. (2012) Systematic identification of genomic markers of drug sensitivity in cancer cells. Nature, 483, 570–575.

3. Yang, W., Soares, J., Greninger, P., Edelman, E.J., Lightfoot, H., Forbes, S., Bindal, N., Beare, D., Smith, J.A., Thompson, I.R., et al. (2013) Genomics of Drug Sensitivity in Cancer (GDSC): a resource for therapeutic biomarker discovery in cancer cells. Nucleic Acids Res., 41, D955–61.

4. Rees, M.G., Seashore-Ludlow, B., Cheah, J.H., Adams, D.J., Price, E.V., Gill, S., Javaid, S., Coletti, M.E., Jones, V.L., Bodycombe, N.E., et al. (2016) Correlating chemical sensitivity and basal gene expression reveals mechanism of action. Nat. Chem. Biol., 12, 109–116.

5. Barretina, J., Caponigro, G., Stransky, N., Venkatesan, K., Margolin, A.A., Kim, S., Wilson, C.J., Lehár, J., Kryukov, G.V., Sonkin, D., et al. (2012) The Cancer Cell Line Encyclopedia enables predictive modelling of anticancer drug sensitivity. Nature, 483, 603–607.

6. Smirnov, P., Kofia, V., Maru, A., Freeman, M., Ho, C., El-Hachem, N., Adam, G.-A., Ba-Alawi, W., Safikhani, Z. and Haibe-Kains, B. (2018) PharmacoDB: an integrative database for mining in vitro anticancer drug screening studies. Nucleic Acids Res., 46, D994–D1002.

7. Reinhold, W.C., Sunshine, M., Liu, H., Varma, S., Kohn, K.W., Morris, J., Doroshow, J. and Pommier, Y. (2012) CellMiner: a web-based suite of genomic and pharmacologic tools to explore transcript and drug patterns in the NCI-60 cell line set. Cancer Res., 72, 3499–3511.

8. Alley, M.C., Scudiero, D.A., Monks, A., Hursey, M.L., Czerwinski, M.J., Fine, D.L., Abbott, B.J., Mayo, J.G., Shoemaker, R.H. and Boyd, M.R. (1988) Feasibility of drug screening with panels of human tumor cell lines using a microculture tetrazolium assay. Cancer Res., 48, 589–601.

9. Boyd, M.R. and Paull, K.D. (1995) Some practical considerations and applications of the national cancer institute in vitro anticancer drug discovery screen. Drug Dev. Res., 34, 91–109.

10. Reinhold, W.C., Varma, S., Sunshine, M., Elloumi, F., Ofori-Atta, K., Lee, S., Trepel, J.B., Meltzer, P.S., Doroshow, J.H. and Pommier, Y. (2019) RNA Sequencing of the NCI-60: Integration into CellMiner and CellMiner CDB. Cancer Res., 79, 3514–3524.

11. Reinhold, W.C., Mergny, J.-L., Liu, H., Ryan, M., Pfister, T.D., Kinders, R., Parchment, R., Doroshow, J., Weinstein, J.N. and Pommier, Y. (2010) Exon array analyses across the NCI-60 reveal potential regulation of TOP1 by transcription pausing at guanosine quartets in the first intron. Cancer Res., 70, 2191–2203.

12. Liu, H., D’Andrade, P., Fulmer-Smentek, S., Lorenzi, P., Kohn, K.W., Weinstein, J.N., Pommier, Y. and Reinhold, W.C. (2010) mRNA and microRNA expression profiles of the NCI-60 integrated with drug activities. Mol. Cancer Ther., 9, 1080–1091.

13. Corsello, S.M., Nagari, R.T., Spangler, R.D., Rossen, J., Kocak, M., Bryan, J.G., Humeidi, R., Peck, D., Wu, X., Tang, A.A., et al. (2020) Discovering the anti-cancer potential of non-oncology drugs by systematic viability profiling. Nat Cancer, 1, 235–248.

14. Iorio, F., Knijnenburg, T.A., Vis, D.J., Bignell, G.R., Menden, M.P., Schubert, M., Aben, N., Gonçalves, E., Barthorpe, S., Lightfoot, H., et al. (2016) A Landscape of Pharmacogenomic Interactions in Cancer. Cell, 10.1016/j.cell.2016.06.017.

15. Haverty, P.M., Lin, E., Tan, J., Yu, Y., Lam, B., Lianoglou, S., Neve, R.M., Martin, S., Settleman, J., Yauch, R.L., et al. (2016) Reproducible pharmacogenomic profiling of cancer cell line panels. Nature, 533, 333–337.

16. Jassal, B., Matthews, L., Viteri, G., Gong, C., Lorente, P., Fabregat, A., Sidiropoulos, K., Cook, J., Gillespie, M., Haw, R., et al. (2020) The reactome pathway knowledgebase. Nucleic Acids Res., 48, D498–D503.

17. Tang, J., Tanoli, Z.-U.-R., Ravikumar, B., Alam, Z., Rebane, A., Vähä-Koskela, M., Peddinti, G., van Adrichem, A.J., Wakkinen, J., Jaiswal, A., et al. (2018) Drug Target Commons: A Community Effort to Build a Consensus Knowledge Base for Drug-Target Interactions. Cell Chem Biol, 25, 224–229.e2.

18. Bairoch, A. (2018) The Cellosaurus, a Cell-Line Knowledge Resource. J. Biomol. Tech., 29, 25–38.

19. Gaulton, A., Hersey, A., Nowotka, M., Bento, A.P., Chambers, J., Mendez, D., Mutowo, P., Atkinson, F., Bellis, L.J., Cibrián-Uhalte, E., et al. (2017) The ChEMBL database in 2017. Nucleic Acids Res., 45, D945–D954.

20. Yu, C., Mannan, A.M., Yvone, G.M., Ross, K.N., Zhang, Y.-L., Marton, M.A., Taylor, B.R., Crenshaw, A., Gould, J.Z., Tamayo, P., et al. (2016) High-throughput identification of genotype-specific cancer vulnerabilities in mixtures of barcoded tumor cell lines. Nat. Biotechnol., 34, 419–423.

21. Picco, G., Chen, E.D., Alonso, L.G., Behan, F.M., Gonçalves, E., Bignell, G., Matchan, A., Fu, B., Banerjee, R., Anderson, E., et al. (2019) Functional linkage of gene fusions to cancer cell fitness assessed by pharmacological and CRISPR-Cas9 screening. Nat. Commun., 10, 2198.

22. Gonçalves, E., Segura-Cabrera, A., Pacini, C., Picco, G., Behan, F.M., Jaaks, P., Coker, E.A., van der Meer, D., Barthorpe, A., Lightfoot, H., et al. (2020) Drug mechanism-of-action discovery through the integration of pharmacological and CRISPR screens. Mol. Syst. Biol., 16, e9405.

23. Kim, S., Chen, J., Cheng, T., Gindulyte, A., He, J., He, S., Li, Q., Shoemaker, B.A., Thiessen, P.A., Yu, B., et al. (2021) PubChem in 2021: new data content and improved web interfaces. Nucleic Acids Res., 49, D1388–D1395.

24. Kim, S., Thiessen, P.A., Cheng, T., Yu, B. and Bolton, E.E. (2018) An update on PUG-REST: RESTful interface for programmatic access to PubChem. Nucleic Acids Res., 46, W563–W570.

25. Kundra, R., Zhang, H., Sheridan, R., Sirintrapun, S.J., Wang, A., Ochoa, A., Wilson, M., Gross, B., Sun, Y., Madupuri, R., et al. (2021) OncoTree: A Cancer Classification System for Precision Oncology. JCO Clin Cancer Inform, 5, 221–230.

26. Mammoliti, A., Smirnov, P., Nakano, M., Safikhani, Z., Eeles, C., Seo, H., Nair, S.K., Mer, A.S., Ho, C., Beri, G., et al. (2021) Orchestrating and sharing large multimodal data for transparent and reproducible research. bioRxiv, 10.1101/2020.09.18.303842.

27. Smirnov, P., Safikhani, Z., El-Hachem, N., Wang, D., She, A., Olsen, C., Freeman, M., Selby, H., Gendoo, D.M.A., Grossmann, P., et al. (2016) PharmacoGx: an R package for analysis of large pharmacogenomic datasets. Bioinformatics, 32, 1244–1246.

28. Volk, M., Staegemann, D., Bosse, S., Häusler, R. and Turowski, K. (2020) Approaching the (big) data science engineering process. In Proceedings of the 5th International Conference on Internet of Things, Big Data and Security. SCITEPRESS - Science and Technology Publications.

29. Hutchinson, B., Smart, A., Hanna, A., Denton, E., Greer, C., Kjartansson, O., Barnes, P. and Mitchell, M. (2021) Towards Accountability for Machine Learning Datasets: Practices from Software Engineering and Infrastructure. In Proceedings of the 2021 ACM Conference on Fairness, Accountability, and Transparency, FAccT ’21. Association for Computing Machinery, New York, NY, USA, pp. 560–575.

30. Karasarides, M., Chiloeches, A., Hayward, R., Niculescu-Duvaz, D., Scanlon, I., Friedlos, F., Ogilvie, L., Hedley, D., Martin, J., Marshall, C.J., et al. (2004) B-RAF is a therapeutic target in melanoma. Oncogene, 23, 6292–6298.

31. Smirnov, P., Smith, I., Safikhani, Z., Ba-alawi, W., Khodakarami, F., Lin, E., Yu, Y., Martin, S., Ortmann, J., Aittokallio, T., et al. (2021) Evaluation of statistical approaches for association testing in noisy drug screening data. arXiv [stat.AP].

32. Press, M.F., Finn, R.S., Cameron, D., Di Leo, A., Geyer, C.E., Villalobos, I.E., Santiago, A., Guzman, R., Gasparyan, A., Ma, Y., et al. (2008) HER-2 gene amplification, HER-2 and epidermal growth factor receptor mRNA and protein expression, and lapatinib efficacy in women with metastatic breast cancer. Clin. Cancer Res., 14, 7861–7870.

33. Xia, W., Mullin, R.J., Keith, B.R., Liu, L.-H., Ma, H., Rusnak, D.W., Owens, G., Alligood, K.J. and Spector, N.L. (2002) Anti-tumor activity of GW572016: a dual tyrosine kinase inhibitor blocks EGF activation of EGFR/erbB2 and downstream Erk1/2 and AKT pathways. Oncogene, 21, 6255–6263.

34. Luna, A., Elloumi, F., Varma, S., Wang, Y., Rajapakse, V.N., Aladjem, M.I., Robert, J., Sander, C., Pommier, Y. and Reinhold, W.C. (2021) CellMiner Cross-Database (CellMinerCDB) version 1.2: Exploration of patient-derived cancer cell line pharmacogenomics. Nucleic Acids Res., 49, D1083–D1093.

