## Supplementary Data for "PharmacoDB 2.0 : Improving scalability and transparency of *in vitro* pharmacogenomics analysis"

### Curation of new datasets

#### 1. NCI-60 Dataset

##### Molecular data compilation

NCI-60 PharmacoSet (PSet) comprises four layers of molecular data including RNA, microRNA, isoform and composite RNA-seq profiles. Given the depth and popularity of NCI-60, there is more than one source of data available for each of the above-mentioned layers. NCI-60 PSet includes the most recent version of the data published for each assay, downloaded directly from CellMiner Processed Data Set section (https://discover.nci.nih.gov/cellminer/loadDownload.do). Robust Multichip Average (RMA) and GeneSpringGX processing methods were chosen for downloading RNA and microRNA data, respectively. Regarding the RNA-seq data, for both isoform and composite profiles, sequencing libraries were generated by TotalScript™ RNA-Seq Kit and sequenced by TruSeq Cluster Kit v3 (Illumina).

Cell line metadata was downloaded directly from CellMiner Cell Line Metadata Section (https://discover.nci.nih.gov/cellminer/celllineMetadata.do) as a text file. Dose response data were available for a total of 163 cell lines including 60 core cell lines, variants of the core cell lines, and additional cell lines listed at the National Cancer Institute website (https://dtp.cancer.gov/discovery_development/nci-60/cell_list.htm). We identified one chinese hamster cell line ("CHO") and two mouse cell lines ("P388" and "P388/ADR") among the additional cell-lines and preserved them in the NCI-60 PSet. Molecular profiles were available only for the 60 core cell lines.

##### Drug curation

NCI-60 metadata references the drugs primarily by National Service Center (NSC) numbers (a total of 55,601 NSC numbers). In NCI-60 PSet curation, it is attempted to map as many NSC numbers as possible to PubChem index names to allow some level of internal standardization to names across datasets. However, PubChem index names could not be fetched for certain NSC numbers using their API. In the absence of PubChem index names, based on availability, NCI-60 drug names from the metadata, or NSC numbers were used respectively as the standard drug names. Each step is explained in further details below.

- NSC numbers were mapped to PubChem index names using our AnnotationGx package (https://github.com/bhklab/AnnotationGx/) functions that hit PubChem PUG-REST API. NSC numbers were first mapped to PubChem compound identifiers (CID) and subsequently, the CIDs were mapped to other drug properties such as SMILES, InChIKeys and PubChem index names.
- For remaining drugs with NSC numbers that could not be mapped to PubChem, NCI-60 drug names from the metadata were used.
- Finally, if an NSC number was not mapped to either Pubchem index names or NCI-60 names, NSC number itself was used as the standard drug identifier.

A flowchart overview of the NCI-60 PSet drug curation process is provided in **Supplementary Figure S1.**

Note -

1. Exceptions such as one to many mapping of NSC to PubChem index names exist. To avoid name repetitions in the dataset, NSC numbers mapping to a unique standard name were concatenated using "///" as the delimiter. For example, the standard name "Virosecurinine" maps to NSC numbers "107413", "107414", and "107415". Accordingly, the corresponding NSC numbers and NCI-60 names were concatenated ("107413///107414///107415", "Securinin///Virosecurinine///Allosecurinin") and mapped to "Virosecurinine" at once.
2. Certain PubChem index names can be long and can include numerous special characters (e.g. "-,(),[]") generating non-human readable names. Such cases were handled using the following internal criteria: if a PubChem index name included more than 10 special characters and if the corresponding NCI-60 name

did not include more than 10 special characters i.e., a human readable name, the latter was used as the standard name (**Supplementary Figure S1)**.

After concatenation of the synonym drugs the PSet included a total of 54774 unique standard names.

The dataset was curated using PharmacoGx (https://bioconductor.org/packages/release/bioc/html/PharmacoGx.html), an R Bioconductor package with a set of functions for storing and analyzing large pharmacogenomics datasets as PSet.

#### Dose response data processing

NCI-60 reports PTC (Percent of Treated cell growth as a fraction of Control cell growth) per log10 of the doses in the dilution series. In NCI-60 PSet, the average of the PTC values is used as the viability measure and the log10 values of the doses are raised to the power of 10 to represent concentrations. Doses are reported in micromolar in the PSet.
In the curation of Drug Dose Response Curves (DDRC), in order to create a unique experiment id, a single experiment was defined as a combination of a drug identifier, (NSC number), a single cell line, and NCI-60 internal experiment ID which account for information such as experiment date and process tracking. For instance, an experiment in which a drug with NSC number "368270" is tested on cell-line "A498" in "renal" cancer with NCI-60 internal experiment ID of "9905NS43", is assigned with the experiment ID of "**368270_A498_Renal_9905NS43**" in the NCI-60 PSet. Further information regarding the components of NCI-60 internal experiment ID is available on NCI wiki (https://wiki.nci.nih.gov/display/NCIDTPdata/NCI-60+Growth+Inhibition+Data).

DDRC parameters such as AAC (area above the curve), IC50 (Half-maximal inhibitory concentration), EC50 (Half-maximal effective concentration), E_inf (drug's maximal effect), and HS (Hill slope) were computed for each experiment using PharmacoGx R package as described previously (1). To obtain mathematically reliable AAC and IC50 values, logistic Hill Curves were fit to  experiments that included at least 4 dose points, using the PharmacoGx package. Some experiments were repeated multiple times at a certain dose introducing technical replicates. Viability measures from such technical replicates were fitted under the same DDRC in NCI-60 PSet to preserve reported data points.

#### 2. PRISM Dataset

##### Molecular data compilation

PRISM PSet does not include molecular profile of the cell lines as PRISM study directly uses cell lines from the Broad-Novartis Cancer Cell Line Encyclopedia (CCLE) project. CCLE molecular profile data is accessible through https://orcestra.ca/ - CCLE_2015 PSet.

##### Drug curation

Synonym drugs in the PRISM dataset were concatenated and mapped to a unique standard name. Overall, the PSet includes 1437 unique standard drug names.

##### Dose response data processing

In PRISM PSet a single experiment is defined as a combination of a drug (distinguished by broad_ids), a single cell line, and screening plates. For example, experiment ID of "**BRD-A00077618-236-07-6**::**HTS002**::**ACH-000007**::**8-bromo-cGMP**" represents an experiment in which a drug with broad_id of "BRD-A00077618-236-07-6" and PRISM drug name of "8-bromo-cGMP" is tested on cell line "ACH-000007" in "HTS002" screening plate.

Same as NCI-60 PSet, Hill Curves were fit to experiments that included at least 4 dose points to achieve mathematically reliable AAC and IC50 measurements. Technical replicates, repetition of a certain experiment at a certain dose, were also observed in PRISM data. Viability values from the technical replicates were fitted under the same DDRCs for the sake of data preservation. PRISM PSet includes recomputed AAC, IC50, EC50, E_inf, and HS values, as well as AUC, IC50 and EC50 values that were published in the PRISM study.

The new PSets, containing both compound response and molecular data for the datasets added in PharmacoDB 2.0 at the time of submitting the manuscript can be accessed at the following DOIs: NCI60 - 10.5281/zenodo.5417785; PRISM - 10.5281/zenodo.5296794; GDSC_v2 - 10.5281/zenodo.3905481; gCSI - 10.5281/zenodo.4737437

### Updated curation of tissues

In the updated process of tissue curation, annotations of cell lines are fetched using data downloaded from Cellosaurus. The “Disease” annotation of a cell line combined with National Cancer Institute thesaurus (NCIt) is the primary information used to annotate tissue. The tissue information from the above resources are matched to the top level of OncoTree hierarchy in two steps - a) if NCIt is present in Cellosaurus and OncoTree, fetch OncoTree tissue name; b) else, map to OncoTree terminology based on best match of disease.

### Web-interface updates

The web application interface has been improved to simplify maintenance and ensure the extensibility of the application with new features in the future. The new client implementation leverages ReactJS as a front-end framework and D3.js and PlotlyJS as libraries for data visualization. Client-server communication is being implemented using GraphQL APIs to ensure greater scalability and standardization. The MySQL database has also been updated with a new schema to accommodate additional drug and cell-line metadata as well as to improve scalability for accumulating pharmacogenomic data.

### Statistical Supplement

As mentioned in the main text, the previous pipeline used to evaluate associations between all feature types and drug response was based around a linear modelling approach. Previously, drug response, measured as the Area Above the Curve value, was modeled as a linear function of tissue specific offsets (arbitrarily taking the alphabetically first tissue as a reference) and the expression, copy number log-ratio, or binary mutation status of each gene independently, as previously described in Smirnov et al. (2). The molecular and drug response values were standardized, which yielded standardized regression coefficients adjusted for tissue specific differences in average response. The significance of the association between the molecular feature and drug response was then tested by comparing the explained variance of the full model to the model containing only the tissue indicator variables, using an analytical F-test. Unfortunately, these standardized coefficients were an unfamiliar measure for most users of PharmacoDB, and they were apt to be misinterpreted as correlation coefficients. Additionally, as mentioned in the text, both drug response and molecular feature distributions are rarely normal, bringing into question the validity of statistical inference based on analytical F-test derived p-values. To address both limitations, we have modified the pipeline to use the (partial) Pearson correlation coefficient as the effect size estimate (adjusted for tissue in the case of pan-cancer analyses). Furthermore, we now augment the results of the analytical test (a t-test in the case of partial or standard Pearson correlation) with the calculation of a permutation test based two-sided p-value. The permutation test is implemented using adaptive stopping with the QUICK-STOP algorithm (3), following the same description as Smirnov, Smith et al. (4). For each drug, tissue and molecular feature type (out of mutation, CNV and RNA expression), the required alpha for rejecting the null hypothesis of no correlation is corrected for multiple testing through a Bonferroni correction on the total number of features of this type tested. It is important to note that the analytical p-value, under violation of the hypothesis, will give a precise but biased estimate of the true p-value, while the early stopping approach to permutation testing gives a less biased (having assumed only independent and identically distributed data) but noisier estimate of the true p-value. The precision of this estimate decreases with distance from the specified alpha level, and as such, PharmacoDB includes both analytical and permutation derived p-values, the former being more useful for accurately sorting associations, while the latter for statistical inference about the significance of a particular correlation between a molecular feature and drug response.

**SUPPLEMENTARY FIGURES**

**Supplementary Figure S1 - A flowchart overview of the NCI-60 PSet compound name curation process.** The NSC number, which is the compound identifier in NCI-60 was mapped to a more human interpretable name in PharmacoDB using the process described in the flowchart. Squares represent entry or exit points from the process, diamonds are Yes/No branches. NCI-60 ID refers to the NCI-60 name when available, NSC number otherwise. NCI-60 IDs were concatenated using “///” in cases where a PubChem index name maps to multiple NCI-60 IDs.

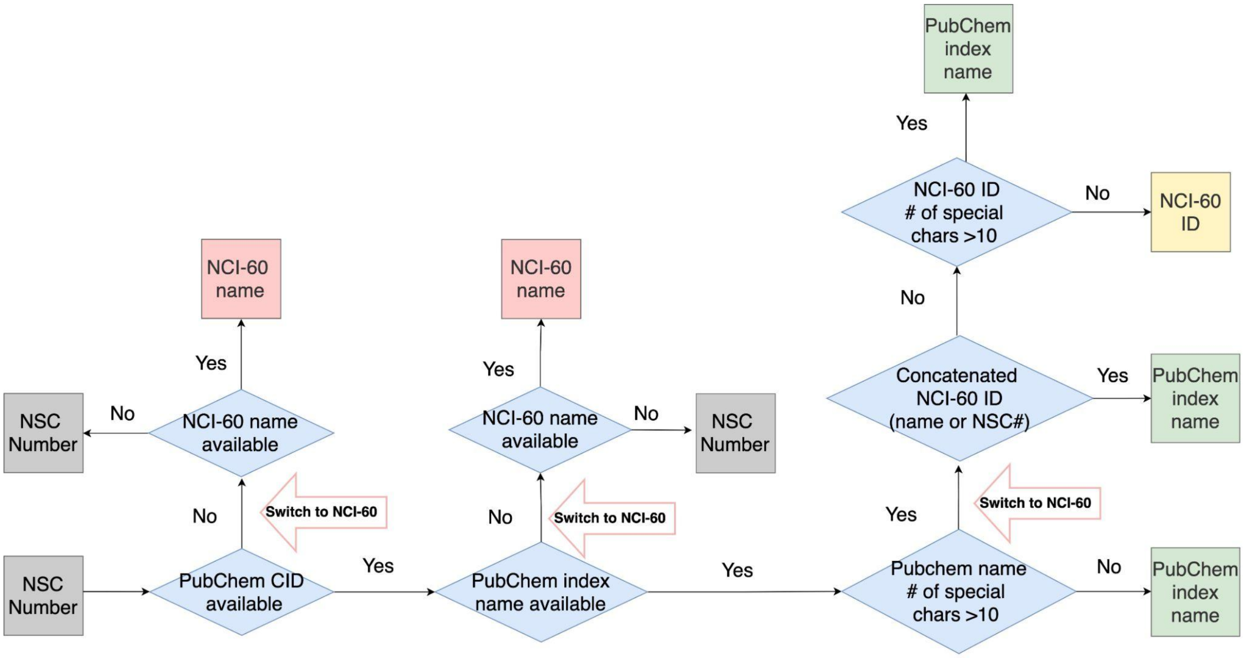


References

1. Smirnov,P., Safikhani,Z., El-Hachem,N., Wang,D., She,A., Olsen,C., Freeman,M., Selby,H., Gendoo,D.M.A., Grossmann,P., *et al.* (2016) PharmacoGx: an R package for analysis of large pharmacogenomic datasets. *Bioinformatics*, **32**, 1244–1246.

2. Smirnov,P., Kofia,V., Maru,A., Freeman,M., Ho,C., El-Hachem,N., Adam,G.-A., Ba-Alawi,W., Safikhani,Z. and Haibe-Kains,B. (2018) PharmacoDB: an integrative database for mining in vitro anticancer drug screening studies. *Nucleic Acids Res.*, **46**, D994–D1002.

3. Hecker,J., Ruczinski,I., Cho,M.H., Silverman,E.K., Coull,B. and Lange,C. (2020) A flexible and nearly optimal sequential testing approach to randomized testing:

QUICK-STOP. *Genet. Epidemiol.*, **44**, 139–147.

4. Smirnov,P., Smith,I., Safikhani,Z., Ba-alawi,W., Khodakarami,F., Lin,E., Yu,Y., Martin,S., Ortmann,J., Aittokallio,T., *et al.* (2021) Evaluation of statistical approaches for association testing in noisy drug screening data. *arXiv [stat.AP]*.
